# SARS-CoV-2 ORF6 disrupts nucleocytoplasmic transport through interactions with Rae1 and Nup98

**DOI:** 10.1101/2020.08.03.234559

**Authors:** Amin Addetia, Nicole A. P. Lieberman, Quynh Phung, Hong Xie, Pavitra Roychoudhury, Lasata Shrestha, Michelle Loprieno, Meei-Li Huang, Keith R. Jerome, Alexander L. Greninger

## Abstract

RNA viruses that replicate in the cytoplasm often disrupt nucleocytoplasmic transport to preferentially translate their own transcripts and prevent host antiviral responses. The *Sarbecovirus* accessory protein ORF6 has previously been shown to be the major inhibitor of interferon production in both SARS-CoV and SARS-CoV-2. SARS-CoV-2 ORF6 was recently shown to co-purify with the host mRNA export factors Rae1 and Nup98. Here, we demonstrate SARS-CoV-2 ORF6 strongly represses protein expression of co-transfected reporter constructs and imprisons host mRNA in the nucleus, which is associated with its ability to co-purify with Rae1 and Nup98. These protein-protein interactions map to the C-terminus of ORF6 and can be abolished by a single amino acid mutation in Met58. Overexpression of Rae1 restores reporter expression in the presence of SARS-CoV-2 ORF6. We further identify an ORF6 mutant containing a 9-amino acid deletion, ORF6 Δ22-30, in multiple SARS-CoV-2 clinical isolates that can still downregulate the expression of a co-transfected reporter and interact with Rae1 and Nup98. SARS-CoV ORF6 also interacts with Rae1 and Nup98. However, SARS-CoV-2 ORF6 more strongly co-purifies with Rae1 and Nup98 and results in significantly reduced expression of reporter proteins compared to SARS-CoV ORF6, a potential mechanism for the delayed symptom onset and pre-symptomatic transmission uniquely associated with the SARS-CoV-2 pandemic.

**Importance:** SARS-CoV-2, the causative agent of COVID-19, is an RNA virus with a large genome that encodes accessory proteins. While these accessory proteins are not required for growth *in vitro*, they can contribute to the pathogenicity of the virus. One of SARS-CoV-2’s accessory proteins, ORF6, was recently shown to co-purify with two host proteins, Rae1 and Nup98, involved in mRNA nuclear export. We demonstrate SARS-CoV-2 ORF6 interaction with these proteins is associated with reduced expression of a reporter protein and accumulation of poly-A mRNA within the nucleus. SARS-CoV ORF6 also shows the same interactions with Rae1 and Nup98. However, SARS-CoV-2 ORF6 more strongly represses reporter expression and co-purifies with Rae1 and Nup98 compared to SARS-CoV ORF6. The ability of SARS-CoV-2 ORF6 to more strongly disrupt nucleocytoplasmic transport than SARS-CoV ORF6 may partially explain critical differences in clinical presentation between the two viruses.

## Introduction

Control over host protein expression allows viruses to suppress the host’s immune response and hijack the host’s translational machinery for expression of viral proteins (1–3). Numerous viruses exert translational control by encoding proteins which target nucleocytoplasmic transport, including the nuclear export of host mRNA (4, 5). The host proteins involved in nucleocytoplasmic transport are targets of multiple viruses including vesicular stomatitis virus (VSV), poliovirus, and Kaposi’s sarcoma-associated herpesvirus (KSHV) (6–8).

Nup98 is a component of the nuclear pore complex and interacts with the RNA export factor Rae1 to bind single stranded RNA and facilitate the translocation of mRNA through the nuclear pore complex (9, 10). The matrix (M) protein of VSV interacts with the Rae1•Nup98 complex at the nucleic acid binding site to prevent single stranded RNA from binding Rae1•Nup98 (6, 11). As a result, mRNA remains trapped within the nucleus and global gene expression within host cells is significantly reduced (6). A single methionine residue surrounded by acidic residues within VSV M is critical for interactions with Rae1 and mutations at this residue impair VSV M’s ability to block mRNA nuclear export (11, 12). ORF10 of KSHV also interacts with Rae1•Nup98 to reduce nuclear export of specific mRNA transcripts (8). Similar to VSV M, ORF10 of KSHV contains a conserved methionine residue surrounded by acidic residues which is likely the interacting motif for the Rae1•Nup98 complex (13).

Severe acute respiratory syndrome coronavirus 2 (SARS-CoV-2), the causative agent of Coronavirus Disease 2019 (COVID-19), is a single stranded RNA virus belonging to the *Betacoronavirus* genus (14). With their large genomes, coronaviruses including SARS-CoV-2 encode accessory proteins that are not required for viral replication, but can contribute the virus’s pathogenicity (15). One of SARS-CoV-2’s accessory proteins, ORF6, was recently shown to co-purify with Rae1 and Nup98 in an affinity purification mass spectrometry screen (13). Similar to VSV M and KSHV ORF10, SARS-CoV-2 ORF6 contains a methionine in its C-terminus that is surrounded by acidic residues, which may facilitate an interaction with the nucleic binding site of the Rae1•Nup98 complex (13). The impact of the putative ORF6-Rae1-Nup98 interactions on export of host mRNA and protein expression has yet to be determined.

In the closely related severe acute respiratory syndrome coronavirus (SARS-CoV), ORF6 was shown to block nuclear import of STAT1 by binding the importin karyopherin alpha 2 (16). SARS-CoV ORF6 has also been shown to downregulate expression of co-transfected constructs (17). However, it is unclear if SARS-CoV ORF6 mediates this repression by restricting mRNA nuclear export via interactions with Rae1 or Nup98.

## Results

### Identification of a 9 amino acid deletion in ORF6 independently arising in multiple clinical SARS-CoV-2 isolates and a serially passaged cultured isolate

While sequencing clinical SARS-CoV-2 isolates, we identified a unique isolate (WA-UW-4752, MT798143) containing a 27-nucleotide, in-frame deletion within ORF6. The isolate was derived from a nasopharyngeal swab (C_T_ 26.1, Hologic Panther Fusion) from an individual who presented to the emergency room after 8 days of fever, cough, and myalgias and was on no specific COVID-19 therapy. The resulting ORF6 protein, referred to as ORF6 Δ22-30, contains a 9-amino acid deletion towards the N-terminus of ORF6 (Figure 1A). The deletion was confirmed by RT-PCR and Sanger sequencing (Figure 1B).

**Figure 1.**
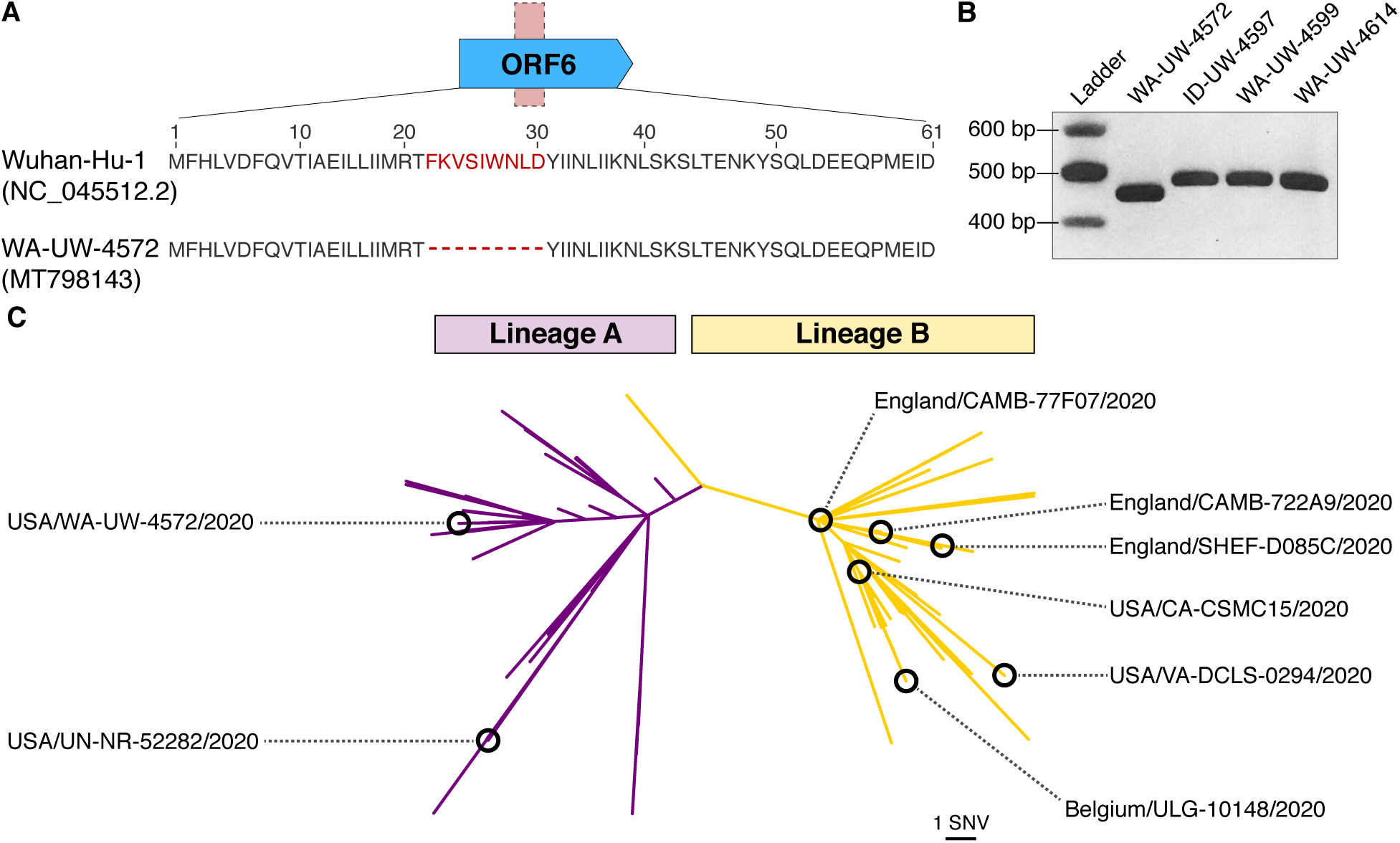
Multiple clinical SARS-CoV-2 isolates and a serially passaged cultured ARS-CoV-2 isolate contain a 9-amino acid deletion in ORF6. A) Schematic presentation of the 9-amino acid deletion in ORF6 identified through whole genome sequencing of the clinical SARS-CoV-2 clinical isolate, WA-UW-4572. B) RT-PCR with primers spanning ORF6 yielded a 452 bp PCR product for WA-UW-4572 rather than a 479 bp product confirming the ORF6 deletion in WA-UW-4572. C) Six other clinical isolates and a cultured isolate with an identified deletion in ORF6 were identified by analyzing the ORF6 sequences of 67,000 publicly available SARS-CoV-2 genomes. The isolates were genetically distinct and belonged to both major SARS-CoV-2 lineages.

We searched for similar ORF6 deletions in over 67,000 SARS-CoV-2 genomes present in GISAID (accessed July 17, 2020) and identified 6 other clinical isolates collected in Virginia, California, Belgium, and the United Kingdom that contained the same ORF6 deletion (Table S1). Additionally, we identified a SARS-CoV-2 isolate with an identical deletion that arose after six serial passages in cell culture (Table S1) (18). Notably, the original clinical isolate used to infect the cells had an intact ORF6 (18). We next performed a phylogenetic analysis (Figure 1C) to determine the genetic relatedness of the 8 strains. The strains differed by 2 – 20 single nucleotide variants and belonged to 4 different lineages, A, A.1, B.1, and B.1.5, as defined by Pangolin (https://github.com/cov-lineages/pangolin) (19), suggesting the ORF6 Δ22-30 deletion arose independently in multiple lineages.

### ORF6 downregulates expression of a co-transfected mCherry reporter

ORF6 putatively interacts with the nuclear export factor Rae1 and the nuclear pore complex component Nup98 (13). VSV M and KSHV ORF10, which both interact with Rae1 and Nup98, downregulate expression of fluorescent or luminescent reporters when co-transfected in cell culture by preventing nuclear export of reporter mRNA (6, 8). We generated a series of N-terminal GFP-tagged ORF6 constructs (Figure 2A) and co-transfected 293T cells with these constructs and a reporter plasmid encoding mCherry. Similar to VSV M, cells expressing the GFP-ORF6 construct showed a significant reduction in mCherry expression (Mean Fluorescent Intensity [MFI]: 0.31; Standard Error [SE]: 0.01; p = 0.01) relative to the cells transfected with GFP alone (MFI: 1.0; SE: 0.09) (Figure 2B-C). Subsequent western blotting and densitometry (Figure 2D-E) further confirmed mCherry expression was downregulated in cells expressing wild-type (WT) ORF6.

**Figure 2.**
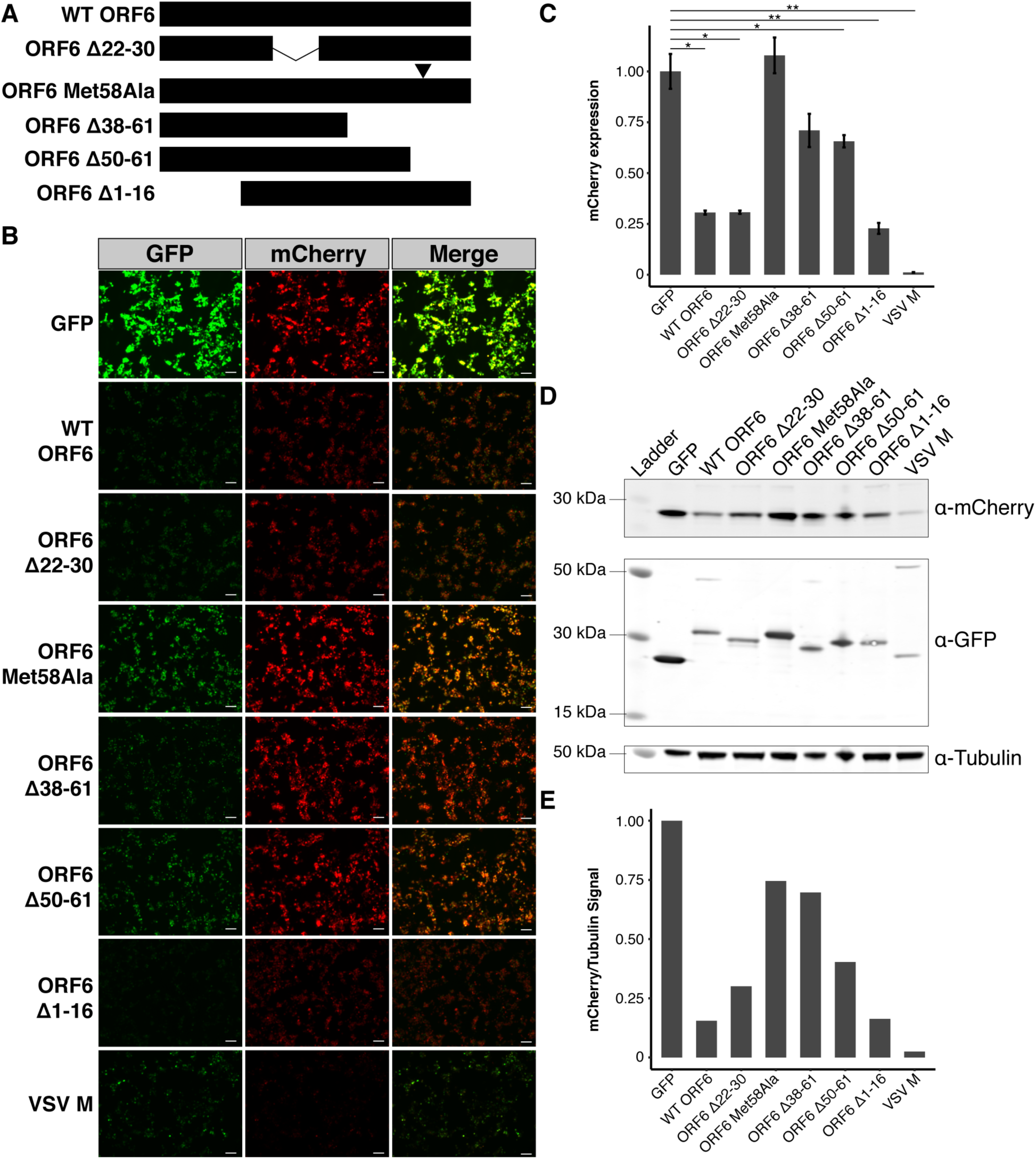
ORF6 of SARS-CoV-2 results in reduced mCherry reporter protein expression in 293T cells. A) Schematic representation of ORF6 constructs used in this study. B) 293T cells were transiently transfected with GFP-tagged constructs and Cherry and visualized 48 hours after transfection. All images were taken with identical fluorescence gain settings. C) The fluorescent intensities for 3 fields per co-transfection were measured with ImageJ and displayed as mean ± standard error. Wild-type (WT) ORF6 caused a significant reduction in mCherry expression. ORF6 constructs with deletions or a single amino acid substitution (Met58Ala) in the C-terminus showed a smaller reduction in mCherry expression than WT ORF6. Cell sates were collected for each of the conditions and D) western blotting and E) densitometry confirmed the role of the C-terminus of ORF6 in reducing protein expression in transfected cells. * p < 0.05; ** p < 0.01. Scale bar: 100 µm.

ORF6 constructs containing deletions in the protein’s N-terminus, ORF6 Δ1-16 (MFI: 0.23, SE: 0.03) and the clinical isolate variant ORF6 Δ22-30 (MFI: 0.31; SE: 0.01), displayed a 3- to 4-fold reduction in mCherry expression (Figure 2B-C) similar to WT ORF6, indicating the N-terminus of ORF6 is not involved in downregulating protein expression. In contrast, mCherry expression was only reduced 1.4- to 1.5-fold in the presence of ORF6 constructs with deletions in the C-terminus, ORF6 Δ38-61 (MFI: 0.71; SE: 0.08) and Δ50-61 (MFI: 0.66; SE: 0.03) (Figure 2B-C).

In VSV M, a motif consisting of a methionine residue surrounded by acidic residues is critical for reducing expression levels of co-transfected reporters. The methionine residue within the motif is conserved between VSV M and KSHV ORF10 and a similar motif with a methionine residue is present in the SARS-CoV-2 ORF6 C-terminus (Figure 1A). We substituted this methionine residue in ORF6 to an alanine, generating the construct ORF6 Met58Ala (Figure 2A). Transfection of ORF6 Met58Ala did not downregulate mCherry expression (MFI: 1.08; SE: 0.09) (Figure 2B-C) suggesting Met58 is critical for the function of ORF6.

### mRNA accumulates in the nucleus in the presence of ORF6

We next investigated whether the reduced mCherry expression observed in the presence of ORF6 was due to the impairment of mRNA nuclear export. We transfected cells with either GFP, GFP-tagged WT ORF6, GFP-tagged ORF6 Met58Ala, or GFP-tagged VSV M and stained the cells with an oligo dT(30) fluorescent probe to visualize mRNA distribution within the transfected cells (Figure 3). In cells transfected with GFP or ORF6 Met58Ala, mRNA was distributed throughout the cell indistinguishable from the mRNA localization pattern in un-transfected cells. In contrast, mRNA in cells expressing WT ORF6 and VSV M was present in multiple foci within the nucleus suggesting the mRNA in these cells was accumulating in the nucleus. The mRNA localization patterns are consistent with the reporter expression (Figure 2B-D) indicating that the downregulation in reporter expression is due to impairment of mRNA nuclear export.

**Figure 3.**
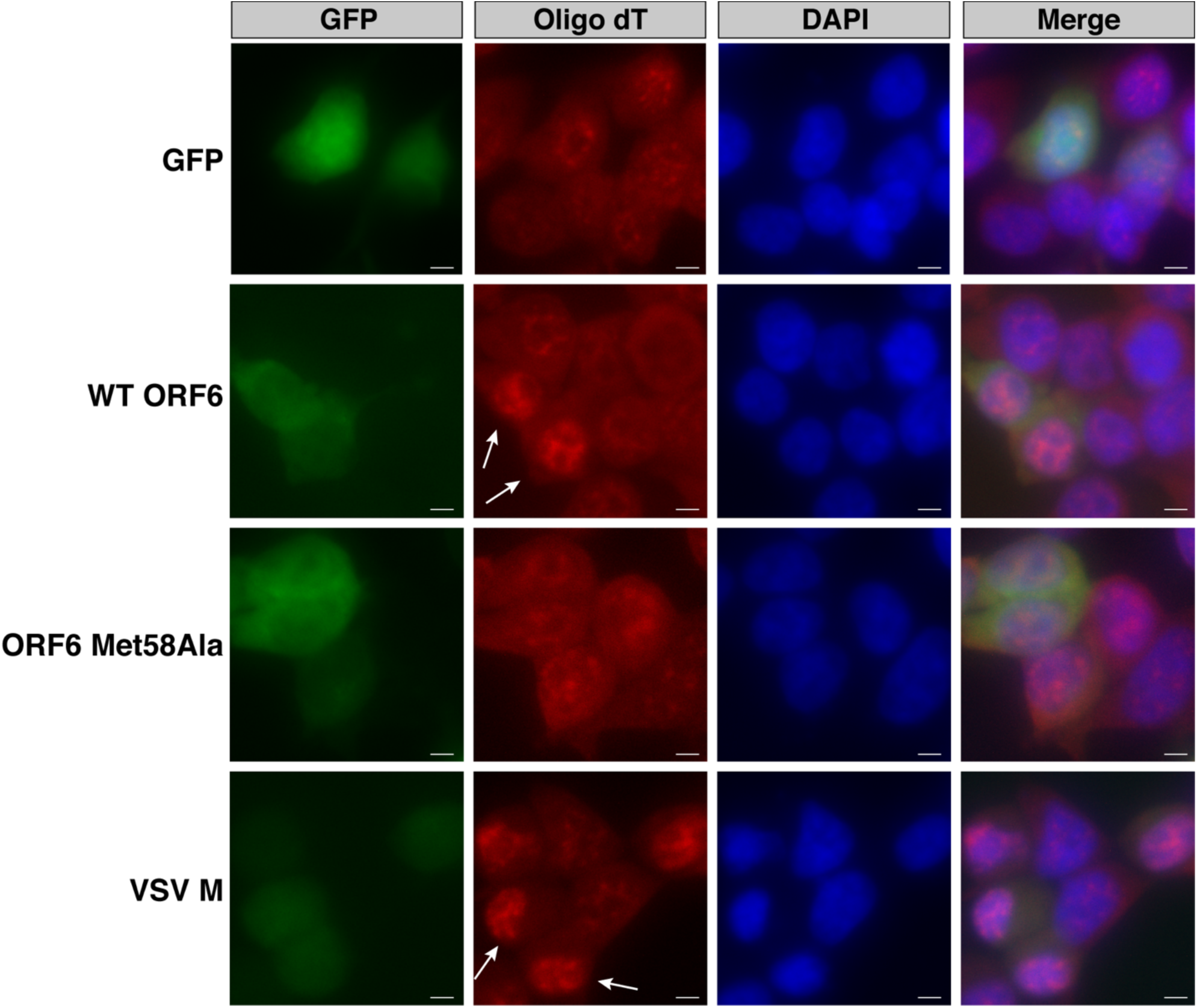
ORF6 causes mRNA to accumulate within the nucleus. 293T cells were transiently transfected with GFP-tagged constructs, incubated for 24 hours, and stained for GFP, poly-A mRNA, and DNA. mRNA in cells transfected with GFP or ORF6 Met58Ala was diffusely present throughout the cell. Cells transfected with ORF6 or VSV M showed an accumulation of mRNA within the nucleus (white arrows). Scale bar: 5 µm.

### The C-terminus of SARS-CoV-2 ORF6 interacts with Rae1 and Nup98

In VSV M and KSHV ORF10, downregulation of co-transfected fluorescent and luminescent reporters and impairment of mRNA nuclear export occurs due to interactions with the nuclear mRNA export factor Rae1 and nuclear pore complex component Nup98 (6, 8). VSV M displaces single stranded RNA in the Rae1•Nup98 complex to prevent nuclear export of host mRNA (11). We hypothesized the inability of the ORF6 C-terminal deletions to downregulate mCherry expression in a similar manner as WT ORF6 (Figure 2B-E) was attributed to the loss of the interaction between these ORF6 constructs and Rae1 and Nup98. We transfected 293T cells with GFP-tagged ORF6 constructs (Figure 2A) and rapidly affinity purified the GFP-tagged proteins. Western blotting on the elutes confirmed WT ORF6, along with ORF6 constructs with N-terminal deletions, interacts with Rae1 and Nup98 (Figure 4). The C-terminal deletion constructs, ORF6 Δ38-61 and ORF6 Δ50-61, did not pull down Rae1 or Nup98 (Figure 4). These data suggest the C-terminus of ORF6 interacts with Rae1 and Nup98, while the N-terminus is not essential for the observed interactions. This is consistent with the observation that C-terminal deletion mutants of ORF6 did not dramatically reduce expression of the mCherry reporter (Figure 2B-E).

**Figure 4.**
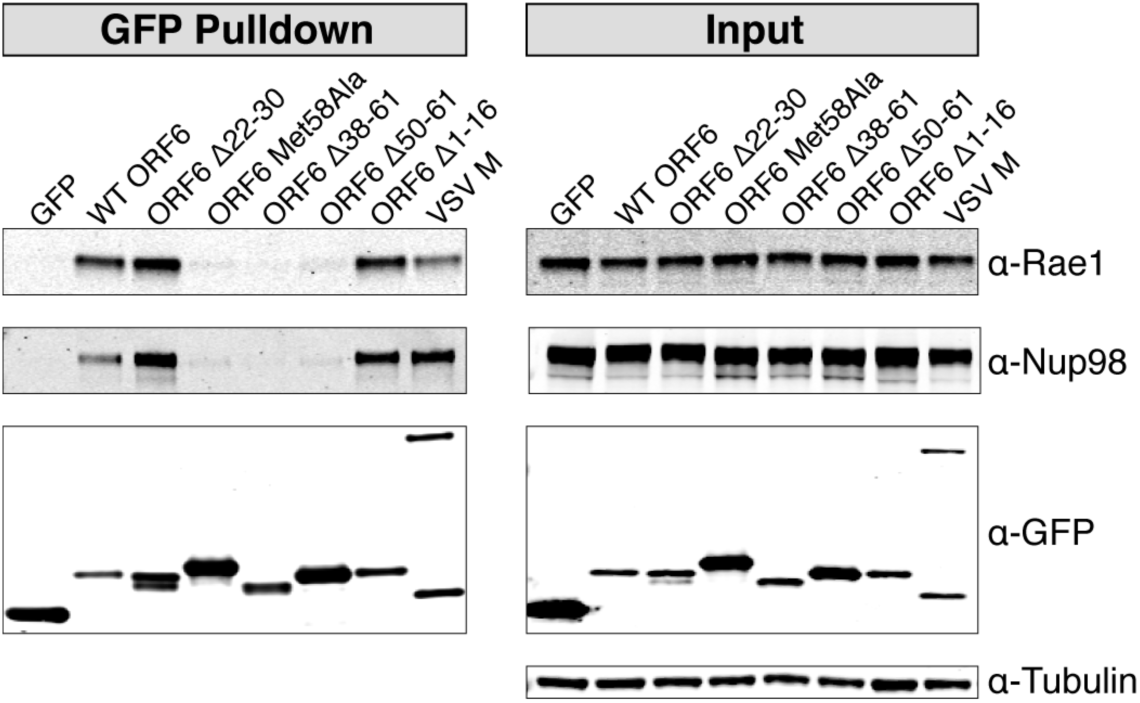
Affinity purification of GFP-tagged constructs. 293T were transiently transfected with GFP-tagged constructs. Forty-eight hours after transfection, the GFP-tagged proteins were rapidly captured using an anti-GFP resin. Western blotting revealed ORF6 interacts with the mRNA nuclear export factor Rae1 and the nuclear ore complex protein Nup98. ORF6 constructs with C-terminal deletions or a substitution did not pull down Rae1 or Nup98.

The methionine residue in the Rae1-Nup98 interacting motif of VSV M forms multiple intermolecular interactions with amino acid residues in the nucleic acid binding site of Rae1 and facilitates the interaction between VSV M and the Rae1•Nup98 complex (11). We hypothesized Met58 of SARS-CoV-2 ORF6 is similarly responsible for interactions with Rae1 and Nup98. Affinity purification of ORF6 Met58Ala revealed it does not interact with Rae1 or Nup98 (Figure 4), confirming the importance of Met58 in the ORF6-Rae1 and ORF6-Nup98 interactions.

### Overexpression of Rae1 restores mCherry reporter expression in cells transfected with ORF6

We next investigated whether we could restore mCherry expression in 293T cells transfected with ORF6 by overexpressing Rae1. Rae1 overexpression restored mCherry expression in a dose-dependent manner (Figure 5A-B). Subsequent western blotting and densitometry confirmed this Rae1 dose-dependent rescue of mCherry expression (Figure 5C-D). These data indicate ORF6’s interaction with Rae1 is responsible for downregulating mCherry reporter expression in cell culture.

**Figure 5.**
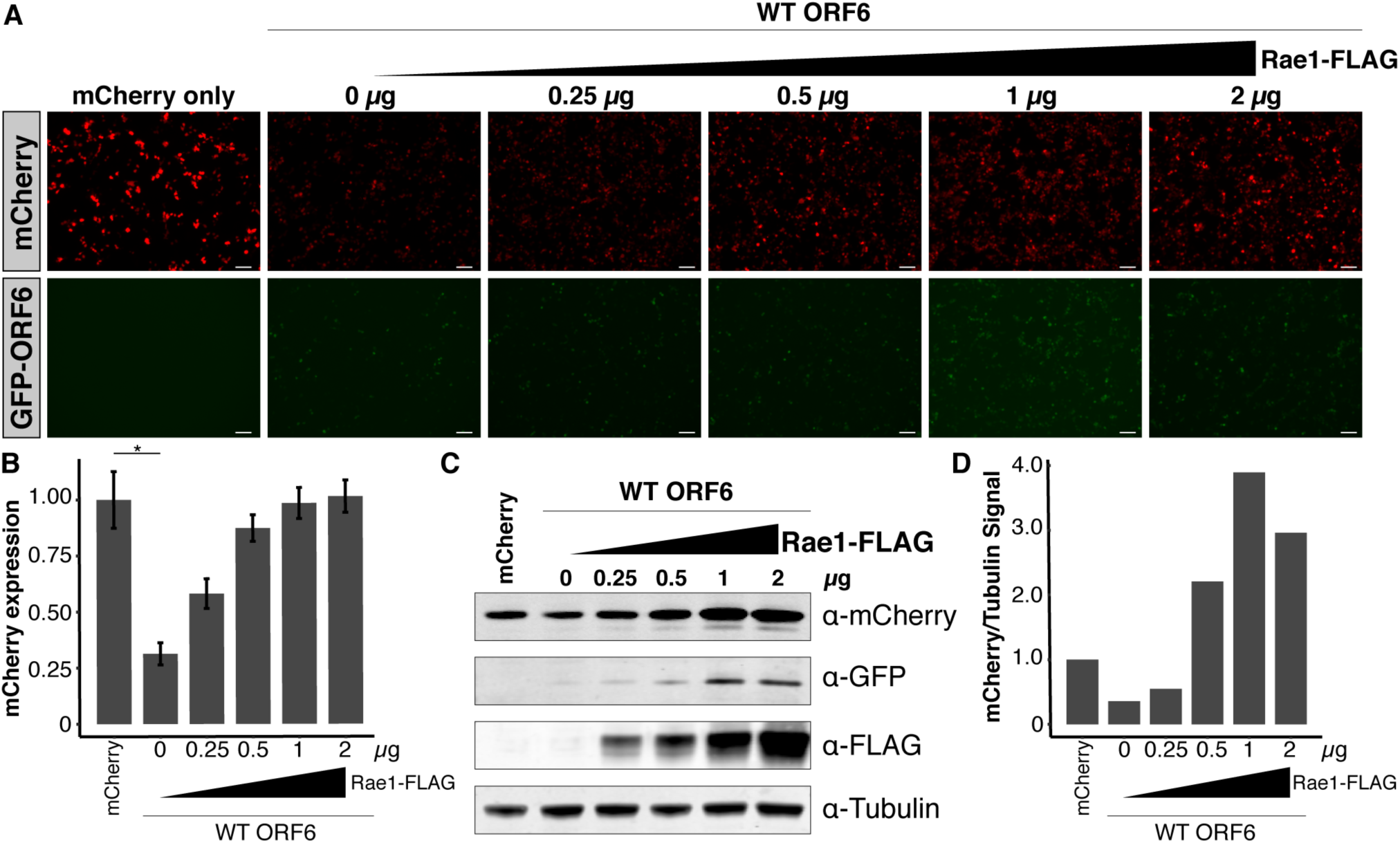
Overexpression of Rae1 rescues mCherry expression in cells transfected with ORF6. A) 293T cells were co-transfected with equal amounts GFP-ORF6 and mCherry and an increasing amount of Rae1. Expression of the fluorescent reporters was visualized and B) quantified 48 hours after transfection. In the presence of ORF6, mCherry expression was restored in a dose-dependent manner. C) Western blotting and D) densitometry confirmed mCherry expression was rescued in a dose-dependent manner with of increasing Rae1-FLAG. * p < 0.05. Scale bar: 100 µm.

### SARS-CoV-2 ORF6 more strongly co-purifies with Rae1 and Nup98 compared to SARS-CoV ORF6

We next compared the relative ability of SARS-CoV ORF6 and SARS-CoV-2 ORF6 to downregulate reporter expression. SARS-CoV ORF6 and SARS-CoV-2 ORF6 share 69% identity by amino acid, including the same methionine residue surrounded by acidic residues (Figure 6A). SARS-CoV ORF6 has been shown to downregulate expression of a co-transfected construct in a dose-dependent manner (19), suggesting its C-terminus may also interact with the Rae1•Nup98 complex.

**Figure 6.**
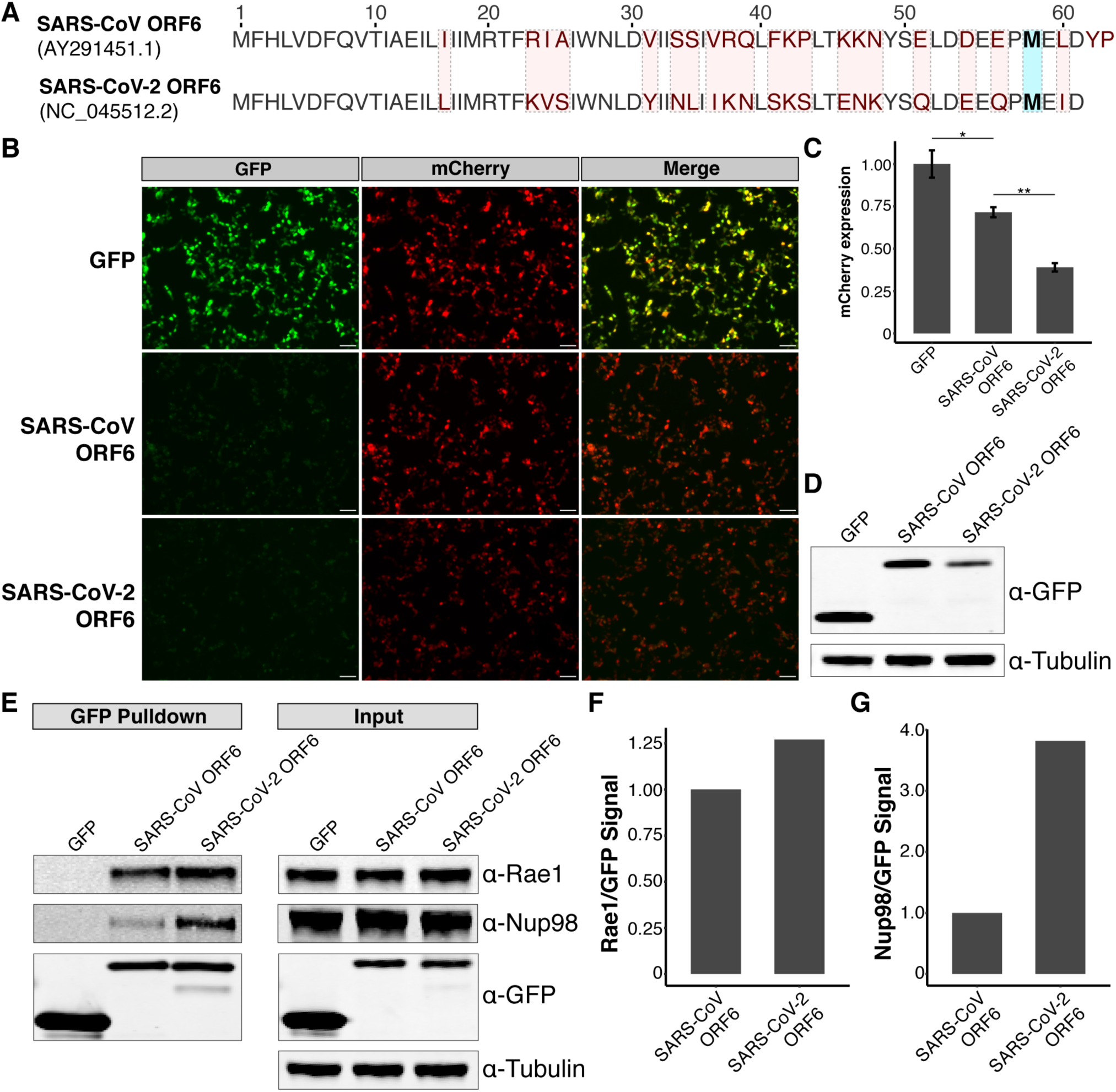
SARS-CoV-2 ORF6 represses reporter expression and copurifies with relatively more Rae1-Nup98 than SARS-CoV ORF6. A) Comparison between the amino acid sequences of ORF6 of SARS-CoV and ORF6 of SARS-CoV-2. Residues differing between the two viruses are highlighted in red. The residue (Met58) implicated in binding the Rae1•Nup98 complex is highlighted in blue. B) 293T were transiently transfected with GFP-tagged constructs and mCherry and visualized 24 hours after transfection. Cells transfected with SARS-CoV-2 ORF6 showed C) significantly reduced mCherry expression compared to those transfected with SARS-CoV ORF6. D) Western blotting showed decreased expression of SARS-CoV-2 ORF6 compared to SARS-CoV ORF6 in 293T cells. E) Affinity purification of GFP-tagged constructs demonstrates both ORF6 of SARS-CoV and ORF6 of SARS-CoV-2 interact with Rae1 and Nup98. Densitometry shows SARS-CoV-2 ORF6 copurifies with relatively more F) Rae1 and G) Nup98 compared SARS-CoV ORF6. * p < 0.05; ** p < 0.01. Scale bar: 100 µm.

We co-transfected 293T cells with GFP-tagged SARS-CoV ORF6 or GFP-tagged SARS-CoV-2 ORF6 and mCherry to assess the impact of these constructs on protein expression. Compared to cells transfected with GFP alone, cells transfected with SARS-CoV ORF6 displayed reduced mCherry expression (MFI: 1; SE: 0.08 vs. MFI: 0.71, SE: 0.03), however this difference was not significant (p = 0.06) (Figure 6B-C). Cells transfected with SARS-CoV-2 ORF6 displayed a significant reduction in mCherry expression compared to cells transfected with SARS-CoV ORF6 (MFI: 0.3; SE: 0.02; p = 0.001) (Figure 6B-C). Western blotting also demonstrated decreased expression of SARS-CoV-2 ORF6 relative to SARS-CoV ORF6, suggesting expression levels do not explain the differential effects on reporter gene expression (Figure 6D).

We hypothesized the differences in mCherry expression between SARS-CoV ORF6 and SARS-CoV-2 ORF6 could be attributed to differences in co-purification of Rae1 and Nup98. We transfected 293T cells with the GFP-tagged constructs and affinity purified the tagged-proteins. Western blotting revealed SARS-CoV ORF6 interacts with Rae1 and Nup98 similar to SARS-CoV-2 ORF6 (Figure 6E). Densitometry on the ratio of prey-to-bait demonstrated SARS-CoV-2 ORF6 co-purified with 1.3-fold more Rae1 (Figure 6F) and 3.8-fold more Nup98 (Figure 6G) compared to SARS-CoV ORF6. These data suggest SARS-CoV-2 ORF6 may more dramatically repress protein expression via a stronger interaction with the Rae1•Nup98 complex compared to SARS-CoV ORF6.

## Discussion

Here, we demonstrate SARS-CoV-2 ORF6 downregulates protein expression and entraps mRNA in the nucleus through interactions with the mRNA nuclear export factor Rae1 and the nuclear pore complex component Nup98. We show reporter repression, mRNA nuclear export, and the host-viral protein-protein interactions are critically dependent on a methionine residue in the ORF6 C-terminus. Additionally, we demonstrate an ORF6 allele with a 9-amino acid deletion which has arisen in multiple clinical SARS-CoV-2 isolates and a serially passaged culture isolate maintains the ability to downregulate expression of co-transfected reporter and interact with Rae1 and Nup98. Finally, we found that SARS-CoV-2 ORF6 more strongly represses reporter expression and more strongly copurifies with Rae1-Nup98 compared to SARS-CoV ORF6.

RNA viruses, including coronaviruses, that replicate in the cytoplasm have mechanisms to suppress cellular translation, which allows these viruses use the host’s translational machinery to preferentially express viral proteins (1–3). In SARS-CoV, ORF6 is not required for growth *in vitro*, however, expression of SARS-CoV ORF6 can increase the replication kinetics of SARS-CoV and the related murine hepatitis virus *in vitro* (20, 21). In addition, recombinant SARS-CoV isolates containing ORF6 grow to higher viral loads than recombinant isolates lacking ORF6 (20). This enhancement in viral growth could be attributed to SARS-CoV ORF6’s ability to downregulate cellular protein expression through interactions with the Rae1•Nup98 complex. As SARS-CoV-2 ORF6 similarly interacts with Rae1 and Nup98, we speculate ORF6 is required for optimal growth of SARS-CoV-2.

In addition to enhancing viral replication, preventing the nuclear export of mRNA can suppress the host’s antiviral response (1–3). The ability of the M protein of VSV to bind Rae1 and Nup98 and prevent mRNA nuclear export is associated with suppressed interferon-ß gene expression (22). Furthermore, VSV strains containing a mutation at the residue responsible for the VSV M-Rae1-Nup98 interactions induce significantly higher interferon-α protein levels than strains containing wild-type alleles of the M protein (23). SARS-CoV-2 ORF6 has been shown to be an interferon antagonist (24) and likely downregulates interferon expression and host antiviral responses in a similar manner to VSV M through its interaction with the Rae1•Nup98 complex.

To date, SARS-CoV-2 has caused several thousand-fold more infections than SARS-CoV in part due to the distinct clinical presentations between the two viruses. COVID-19 patients display peak viral loads and maximum infectivity upon the onset of symptoms rather than after the onset of symptoms which is typical in patient with SARS (25). Furthermore, asymptomatic transmission was infrequently reported for SARS-CoV (26, 27), however, pre-symptomatic and asymptomatic transmission have been a defining challenge of the current SARS-CoV-2 pandemic (28–30). Both the delayed onset of clinical symptoms and pre-symptomatic and asymptomatic transmission of SARS-CoV-2 could be attributed to increased potency of interferon antagonization in SARS-CoV-2 compared to SARS-CoV. ORF6 has already been shown to be a major interferon antagonist in both SARS-CoV and SARS-CoV-2 (16, 24, 31). ORF6 is one of the least similar accessory proteins (69% identical by amino acid) between the two viruses. Coupled with our demonstration of SARS-CoV-2 ORF6 more strongly downregulating protein expression and co-purifying with more Rae1 and Nup98 than SARS-CoV ORF6, the differences between SARS-CoV ORF6 and SARS-CoV-2 ORF6 could explain at least some of the differences in clinical presentations between SARS and COVID-19.

Large-scale SARS-CoV-2 genomic surveillance projects have demonstrated deletions can arise within the accessory genes of SARS-CoV-2 (32–34). Notably, none of these deletions have arisen in multiple SARS-CoV-2 lineages through multiple independent genomic rearrangement events. Our identification of 7 unrelated clinical isolates with the same ORF6 deletion suggests this deletion may be repeatedly selected for in SARS-CoV-2. This is further evidenced by the identification of a cultured SARS-CoV-2 that acquired the same deletion after successive passages in Vero cells (18). Similar to wild-type ORF6 allele, the clinical allele, ORF6 Δ22-30, can repress expression of a co-transfected reporter and still retains the Rae1•Nup98 interacting motif of ORF6. Further work is required to understand the functional role of the ORF6 N-terminus and determine the selective pressures which are repeatedly selecting for the observed deletion.

Our study has a number of limitations. We relied on cellular overexpression systems of ORF6 which rely on nuclear export, making study of a likely nuclear export inhibition factor difficult. As such, our results may not perfectly reflect the degree to which host mRNA nuclear export and protein expression is downregulated during SARS-CoV-2 infection, which does not rely on nuclear export for ORF6 expression.

More comparative work between SARS-CoV-2 and SARS-CoV ORF6 is needed in the context of viral replication. It would be intriguing to swap ORF6 between SARS-CoV and SARS-CoV-2 isolates to test the hypothesis that ORF6 is the major determinant of interferon antagonization and delayed symptom onset in animal models of SARS-CoV-2. Furthermore, additional work is required to understand the degree of mRNA export inhibition in both viruses.

In summary, our results demonstrate the accessory protein ORF6 of SARS-CoV-2 strongly inhibits reporter protein expression and imprisons mRNA in the nucleus via its interactions with the mRNA nuclear export factor Rae1 and the nuclear pore complex component Nup98. As ORF6 is a major interferon antagonist in *Sarbecoviruses*, differences in ORF6 sequence content may be major determinants of differences in clinical presentation among these viruses that so clearly have the world’s attention.

## Methods

### Specimen collection and whole genome sequencing of SARS-CoV-2 positive clinical specimens

Whole genome sequencing of SARS-CoV-2 positive clinical specimens was conducted as part of an ongoing University of Washington Institutional Review Board-approved study (STUDY00000408) (35–38). Nasopharyngeal swabs were collected from patients suspected to have an infection with SARS-CoV-2 and stored in 3 mL of viral transport medium. RNA was extracted from 140 µL of medium using the Qiagen Biorobot. Sequencing libraries were prepared as previously described (32). Briefly, RNA was treated with TURBO DNase (Thermo Fisher) and first-strand cDNA was synthesized using Superscript IV (Thermo Fisher) and random hexamers (IDT). Double stranded cDNA was created using Sequenase Version 2.0 (Thermo Fisher) and purified using 1.6x volumes of AMPure XP beads (Beckman-Coulter). Multiplex amplicon sequencing libraries were constructed using Swift Biosciences’ SARS-CoV-2 Multiplex Primer Pool and Normalase Amplicon kit and sequenced on a 2 × 300 bp run on an Illumina MiSeq.

712,394 sequencing reads were obtained for the clinical SARS-CoV-2 sample, WA-UW-4752. Sequencing reads were quality- and adapter-trimmed using Trimmomatic v0.38 (ILLUMINACLIP:TruSeq3-PE-SNAP.fa:2:30:10:1:true LEADING:3 TRAILING:3 SLIDINGWINDOW:4:30 MINLEN:75) (39) and aligned to the SARS-CoV-2 reference genome (NC_045512.2) using BBMap version 38.70 (sourceforge.net/projects/bbmap/). Sequence reads were then clipped of synthetic PCR primers using Primerclip (https://github.com/swiftbiosciences/primerclip) and the final sequence alignment was visualized in Geneious version 11.1.4 (40).

The deletion identified within ORF6 of WA-UW-4752 was confirmed by reverse transcription PCR and Sanger sequencing. For reverse transcription, single-stranded cDNA was constructed using Superscript IV. The resulting cDNA was used as template for PCR with Phusion High-Fidelity Polymerase (Thermo Fisher) and the following primers: 5’ ATCACGAACGCTTTCTTATTAC 3’ and 5’ CTCGTATGTTCCAGAAGAGC 3’. PCR was conducted using the following conditions: 98°C for 30 seconds followed by 35 cycles of 98°C for 10 seconds, 55°C for 15 seconds, and 72°C for 30 seconds followed by a final extension at 72°C for 5 minutes. The resulting amplicons were run on a 2% agarose gel, extracted from the gel using the QIAquick Gel Extraction kit (Qiagen), and Sanger sequenced by Genewiz, Inc. with the same primers used for PCR.

Other strains with the same deletion in ORF6 were identified by querying GISAID (accessed July 17, 2020). The genetic relatedness of these strains was assessed by aligning the genomes of these strains as well as 110 other global clinical SAR-CoV-2 strains using MAFFT v7.453 (41). A phylogenetic tree was generated using RAxML version 8.2.11 (42) and visualized with R (version 3.6.1) using the ggtree package (43). Strains were further classified using the web-based lineage assigner, Pangolin (https://pangolin.cog-uk.io/) (19).

### Cloning

The wild-type, N- and C-terminal mutant SARS-CoV-2 ORF6 constructs were amplified from double-stranded cDNA from a previously sequenced clinical SARS-CoV-2 isolate (WA12-UW8; EPI_ISL_413563) using the primers listed in Table S2. CloneAmp Hi-Fi PCR Premix (Takara) and the following PCR conditions were used to generate the amplicons: 98°C for 2 minutes followed by 35 cycles of 98°C for 10 seconds, 55°C for 15 seconds, and 72°C for 30 seconds followed by a final extension for 72°C for 5 minutes. ORF6 Δ22-30 was amplified from WA-UW-4572 (MT798143) and the matrix protein from vesicular stomatitis virus was amplified from pVSV eGFP dG (a gift from Connie Cepko; Addgene plasmid #31842) as described above using the primers listed in Table S2. A gBlock gene fragment (IDT) for ORF6 of SARS-CoV was synthesized based on the genome sequence of SARS-CoV isolate TW1 (AY291451.1). The resulting amplicons and gene fragment were then cloned into a modified pLenti CMV GFP Puro plasmid (a gift from Eric Campeau & Paul Kaufman; Addgene plasmid #17448), which contains a 3’ WPRE sequence following the insert and a 3’ SV40 polyadenylation signal after the puromycin resistance cassette, with an N-terminal GFP tag using the In-Fusion HD Cloning kit (Takara).

For cloning of Rae1, RNA was extracted from 239T cells using the RNeasy Miniprep kit (Qiagen) and cDNA was synthesized using Superscript IV and oligo dT (IDT). Rae1 was then amplified from the resulting cDNA using the primers listed in Table S2 and CloneAmp Hi-Fi PCR Premix under the following PCR conditions: 98°C for 2 minutes followed by 35 cycles of 98°C for 10 seconds, 55°C for 15 seconds, and 72°C for 1 minute followed by a final extension for 72°C for 5 minutes. The resulting amplicon was cloned into a modified pcDNA4-TO vector with a C-terminal FLAG tag using the In-Fusion HD Cloning kit.

### Cell culture and ORF6-mCherry transient co-transfections

293T cells were maintained in Dulbecco’s Modified Eagle’s Medium (GE Healthcare Life Sciences) supplemented with 10% FBS (Sigma-Aldrich), 1x HEPES (Thermo Fisher), and 1x GlutaMAX (Thermo Fisher) (293T media). Transient co-transfections with GFP-tagged constructs and a modified pLenti CMV Puro vector encoding the fluorescent reporter mCherry were conducted in 6-well plates. The day prior to transfection, 500,000 293T cells were plated into each well of the 6-well plate and grown overnight until they reached approximately 50% confluency. The cells were then transfected with 2 µg of GFP-tagged construct and 2 µg of mCherry using a 3:1 ratio of PEI MAX (Polysciences) in Opti-MEM (Thermo Fisher). Cells were incubated for 24-48 hours following transfection and visualized using the EVOS M5000 Imaging System (Thermo Fisher) with GFP and Texas Red filter cubes.

mCherry fluorescence intensities were measured with ImageJ v1.53a by an individual blinded to experimental design. All images were 8-bit grayscale and 2048×1536 (3.1 megapixels). Background thresholds were set at the same level across all images, and mean fluorescence intensity of regions of interest greater than 200 pixels calculated. Three fields were analyzed for each experimental condition. The mean fluorescent intensity for each field was calculated after adjusting for background fluorescence signal and normalized to the control condition. Difference in mean fluorescent intensities between experimental conditions were assessed in R using the unpaired t-test.

Cell lysates were collected 24-48 hours after transfection using RIPA buffer (Thermo Fisher). The total protein content was measured using the Pierce BCA Protein Assay kit (Thermo Fisher) and 7.5 µg of lysate was run on a 4-12% Bis-Tris sodium dodecyl sulfate (SDS)-polyacrylamide gel with MOPS running buffer (Invitrogen) under reducing conditions. The samples were then transferred to a 0.45 μm nitrocellulose membrane using the XCell Blot II module (Invitrogen). Blotting was performed using the following primary antibodies: 1:1,000 anti-GFP (Cell Signaling; clone 4B10), 1:500 anti-mCherry (Cell Signaling; clone E5D8F), and 1:1,000 anti-alpha Tubulin (Cell Signaling; clone DM1A), which was followed by staining with either 1:10,000 IRDye 680RD anti-Mouse IgG secondary antibody (Licor) or 1:5,000 IRDye 800CW anti-Rabbit IgG secondary antibody (Licor). Blots were then visualized on a Licor Odyssey imager using Image Studio version 2.0.

### Oligo dT in situ hybridization

293T cells were plated in 48-well plates at a density of 60,000 cells per well and grown overnight to approximately 50% confluency. Cells were transfected with 200 ng of plasmid DNA as described above and incubated for 24 hours. The cells were then washed with PBS (pH 7.4; without Ca^2+^ or Mg^2+^) (Thermo Fisher) and fixed with 4% paraformaldehyde. The fixed cells were permeabilized with methanol and rehydrated in 70% ethanol followed by 1M Tris-HCl (pH 8.0) (Invitrogen). The monolayer was then covered with hybridization buffer (1 mg/mL yeast tRNA, 0.005% bovine serum, 10% dextran sulfate and 25% formamide in 2x SSC buffer) containing an oligo dT(30) probe with an Alexa Fluor 594 fluorophore (IDT) attached to the 5’ end of the probe and incubate overnight at 37°C. The hybridization buffer was removed and the cells were washed once with warmed 4x SSC buffer (Thermo Fisher), once with warmed 2x SSC buffer, and twice with room temperature 2x SSC buffer.

The cells were then blocked with 1% bovine serum in PBS containing 0.1% Tween 20 (PBST) for 1 hour. To detect the GFP-tagged proteins, the cells were incubated with a FITC conjugated anti-GFP antibody (Abcam) for 1 hour. The antibody was removed and the cells were washed three times with PBST. Nuclear staining was completed by incubating the cells in a 300 nM DAPI solution (Thermo Fisher) for 5 minutes. The cells were washed twice with PBS and visualized on an EVOS M5000 with the GFP, Texas Red, and DAPI filter cubes.

### Affinity Purification of GFP-tagged constructs

The day prior to transient transfection, 10-cm plates were seeded with 4×10^6^ 293T cells and grown overnight to approximately 50% confluency. The cells were transfected with 7 µg of plasmid DNA using a 3:1 ratio of PEI MAX in Opti-MEM. Forty-four to 48 hours after transfection, the cells were washed with PBS and collected using PBS containing 0.1 mM EDTA. The cells were pelleted, resuspended in 500 μL TEN (50mM Tris 8.0, 150mM NaCl, and 1mM EDTA) buffer with 0.5% NP-40, and lysed by rotation for 45-60 minutes at 4°C. The lysates were centrifuged at 13,000 RPM for 5 minutes at 4°C and the supernatant was transferred to a new tube and cleared of residual IgG by rotation with Protein G Sepharose 4 Fast Flow (GE Healthcare Life Sciences) for 30 minutes at 4°C. Cleared lysates were transferred to new tubes and incubated overnight at 4°C with anti-GFP Nanobody Affinity gel (BioLegend). The affinity gel was then pelleted and washed twice using TEN buffer with 0.1% NP-40 and resuspended in equal volumes of NuPage LDS Sample Buffer (Thermo) containing 143 mM 2-Mercaptoethanol (Sigma-Aldrich). Western blotting using the elutes from affinity purification and the pre-purified input lysates were performed as described above with the following primary antibodies: 1:1,000 anti-GFP, 1:1,000 anti-alpha Tubulin, 1:2,000 anti-Rae1 (Abcam; clone EPR6923) and 1:1,000 anti-Nup98 (Abcam; clone 2H10).

### Rae1 rescue of mCherry expression

293T cells were plated in 6-well plates at a seeding density of 500,000 cells per well and grown overnight until they reached approximately 50% confluency. Cells were then transfected with 0.5 µg of the GFP-SARS-CoV-2 wild-type ORF6 construct, 0.5 µg of mCherry, and 0, 0.25, 0.5, 1, or 2 µg of Rae1-FLAG using a 3:1 ratio of PEI MAX in Opti-MEM. GFP and mCherry expression were visualized 44-48 hours following transfection using the EVOS M5000 imaging system with GFP and Texas Red filter cubes. Western blotting was performed as described above with the following primary antibodies: 1:1,000 anti-GFP, 1:500 anti-mCherry, 1:1,000 anti-alpha Tubulin, and 1:1,000 anti-FLAG (Sigma; clone M2).

### Data Availability

Sequencing reads and genome assemblies are available under NCBI BioProject PRJNA610428.

