## Supplemental Tables for "SARS-CoV-2 ORF6 disrupts nucleocytoplasmic transport through interactions with Rae1 and Nup98"

| **Strain Name** | **Lineage** | **Collection Location** | **Accession Number** |
| --- | --- | --- | --- |
| Belgium/ULG-10148/2020 | B.1 | Belgium | EPI_ISL_447145 |
| England/CAMB-722A9/2020 | B.1 | England, UK | EPI_ISL_439593 |
| England/CAMB-77F07/2020 | B.1 | England, UK | EPI_ISL_441819 |
| England/SHEF-D085C/2020 | B.1.5 | England, UK | EPI_ISL_475487 |
| USA/CA-CSMC15/2020 | B.1 | California, USA | EPI_ISL_475624 |
| USA/UN-NR-52282/2020 | A | Cell culture isolate | EPI_ISL_456656 |
| USA/VA-DCLS-0294/2020 | B.1 | Virginia, USA | EPI_ISL_463097 |
| USA/WA-UW-4572/2020 | A.1 | Washington, USA | MT798143 |

**Table S1.** Clinical and cultured SARS-CoV-2 isolates with a 9 amino acid deletion in ORF6 identified by analyzing ORF6 sequences from over 67,000 SARS-CoV-2 strains (<https://www.gisaid.org/>; accessed July 17, 2020).

**Table S2.** Primers used in this study.

| **Name** | **Sequence (5´ to 3´)** |
| --- | --- |
| ORF6-pLenti-GFP-F | GCAGCGGTGGCGGTGGATCCATGTTTCATCTCGTTGACTTTC |
| ORF6-pLenti-GFP-R | TCCAGAGGTTGATTGTCGACGCGCCCGGGTTAATCAATCTCCATTGGTTGC |
| ORF6-M58A-pLenti-GFP-R | TCCAGAGGTTGATTGTCGACGCGCCCGGGTTAATCAATCTCCGCTGGTTGCTCTTCAT |
| ORF6-del38-61-pLenti-GFP-R | TCCAGAGGTTGATTGTCGACGCGCCCGGGTTAAATTATGAGGTTTATGATGTAATC |
| ORF6-del50-61-pLenti-GFP-R | TCCAGAGGTTGATTGTCGACGCGCCCGGGTTAATATTTATTCTCAGTTAGTGAC |
| ORF6-del1-16-pLenti-GFP-R | GCAGCGGTGGCGGTGGATCCATTATTATGAGGACTTTTAAAG |
| VSV-M-pLenti-GFP-F | GCAGCGGTGGCGGTGGATCCATGAGTTCCTTAAAGAAGATTCTC |
| VSV-M-pLenti-GFP-R | TCCAGAGGTTGATTGTCGACGCGCCCGGGTCATTTGAAGTGGCTGATAGAATCC |
| Rae1-pCDNA4TO-FLAG-F | TACCGAGCTCGGATCCATGAGCCTGTTTGGAACAACC |
| Rae1-pCDNA4TO-FLAG-R | CACCGCCTCCCTCGAGCTTCTTATTCCTGGGCTTTAGC |
